# Squidly: Enzyme Catalytic Residue Prediction Harnessing a Biology-Informed Contrastive Learning Framework

**DOI:** 10.1101/2025.06.13.659624

**Authors:** William JF Rieger, Mikael Boden, Frances Arnold, Ariane Mora

**Affiliations:** School of Chemistry and Molecular Biology, University of Queensland, Cooper Road, 4072, Queensland, Australia; Division of Chemistry and Chemical Engineering, California Institute of Technology, California Blvd., 91125, California, USA; AITHYRA GmbH, Research Institute for Biomedical Artificial Intelligence of the Austrian Academy of Sciences, Helmut-Qualtinger-Gasse 2, Stg. 2, A-1030 Vienna, Austria

## Abstract

Enzymes present a sustainable alternative to traditional chemical industries, drug synthesis, and bioremediation applications. Because catalytic residues are the key amino acids that drive enzyme function, their accurate prediction facilitates enzyme function prediction. Sequence similarity-based approaches such as BLAST are fast but require previously annotated homologs. Machine learning approaches aim to overcome this limitation; however, current gold-standard machine learning (ML)-based methods require high-quality 3D structures limiting their application to large datasets. To address these challenges, we developed Squidly, a sequence-only tool that leverages contrastive representation learning with a biology-informed, rationally designed pairing scheme to distinguish catalytic from non-catalytic residues using per-token Protein Language Model embeddings. Squidly surpasses state-of-the-art ML annotation methods in catalytic residue prediction while remaining sufficiently fast to enable wide-scale screening of databases. We ensemble Squidly with BLAST to provide an efficient tool that annotates catalytic residues with high precision and recall for both in- and out-of-distribution sequences.

## Introduction

Enzymes are efficient, non-toxic and sustainable alternatives to traditional chemical catalysts in applications such as bio-remediation, pharmaceutical and biofuel production^1^. Catalytic residues refer to the amino acids in the protein sequence directly involved in the mechanism of action. Catalytic residues are involved both directly and indirectly in catalysis; for example, they can stabilize transition states ^2^, act as direct electron donors, or interact with secondary residues, water molecules, or cofactors that are then directly involved in the catalytic mechanism. As catalytic activity depends on specific structural conformations and sequence context, identifying catalytic residues remains a challenge.

Catalytic residue annotations can be used to infer protein function prediction and enable insights into mechanisms of action and evolutionary relationships ^3–5^. Annotations are used to facilitate enzyme engineering methods such as directed evolution and rational design ^6^, and can help construct the theoretical catalytic sites known as theozymes that are used to condition *de novo* enzyme design ^7^. Experimental methods to determine catalytic residues are time-consuming, labour-intensive, and costly, as they often require experimentally determined enzyme-substrate structures. To expedite annotation, computational approaches have been developed to predict catalytic residues from protein sequences or predicted structures ^8–11^, using annotations in databases such as UniProt ^12^ and M-CSA, a manually curated enzyme mechanism database ^13^. Existing computational methods can be broadly grouped into three categories: 1) similarity-based methods and sequence alignment, 2) structural methods, or 3) statistical and machine learning approaches.

Similarity-based approaches are effective when homologous sequences have been previously annotated. Structure-based approaches use a variety of geometric search algorithms to find potential binding pockets and then map sequence conservation scores to these regions to predict catalytic residues with higher specificity ^14–16^. Geometric methods improve the accuracy of approaches reliant on homology, while the limitation of requiring homologous sequences remains. Data-driven approaches such as statistical models or machine learning algorithms offer a more flexible framework ^9,11,17^ but are restricted by the quality and diversity of datasets available for training. The current machine learning (ML) state-of-the-art (SOTA) tools apply multimodal deep learning strategies and often emphasise the value of the structural modality. Shen et al. ^10^ combined an adaptive edge-gated graph attention neural network (AEGAN) with graphs constructed using the 3D structure and sequence-derived positional-specific scoring matrices. SCREEN leverages protein sequence and structure by combining graphical convolutional neural networks and contrastive learning on Protein Language Models (PLMs) ^9^. EasIFA, developed by Wang et al. ^11^, predicts both catalytic residues and binding sites by incorporating graph attention representations of chemical reactions with PLM information and structure representations. Despite the reported improvements on sequence-based methods by multi-modal approaches, current benchmarks inherently obscure the reliance on structural prediction as they often contain sequences with known structures, which are more likely to be predicted accurately by tools such as AlphaFold ^18^. This bias may limit the evaluation of generalisability to unseen data, i.e. enzymes without experimentally derived structures. Furthermore, computing structures remains computationally intensive.

In comparison to structural models PLMs learn the distribution of amino acids across protein sequences. PLMs embed biologically meaningful representations that capture complex dependencies between residues within a protein sequence ^19–21^. These models are trained in an unsupervised manner on large numbers (billions) of diverse sequences ^22,23^. Remote homology search and alignment algorithms based on PLM embeddings have been reported to outperform sequence similarity-based methods at low sequence identities ^24,25^. In enzyme function prediction, models highlight the utility of PLM embeddings in combination with contrastive learning, to achieve SOTA ML performance in the prediction of enzyme function (by proxy of enzyme classification numbers) and associated reactions catalysed ^26–28^; however, similarity-based approaches (e.g. BLAST or reaction distance) perform comparably ^28^. We hypothesised that using PLM embeddings to predict catalytic residues would complement sequence similarity approaches in low homology settings.

Contrastive learning is a discriminative modelling approach that facilitates learning by distinguishing between paired observations in data that are “similar” or “different”, thus enabling information to be extracted from dense PLM embeddings ^29^. Data engineering, such as choosing a dataset to enhance specific relationships ^30^, enables domain expertise to inform pair schemes for training. For example, it is common to explicitly incorporate and maximise “hard negatives” (samples that are dissimilar but close in the latent space) to refine the model’s ability to discriminate subtle differences in the feature space ^31–33^.

Although pair schemes can be generated automatically in contrastive learning frameworks ^30^, we posited that the physical principles of enzyme functions could be used to maximise hard negatives and create an accurate catalytic residue prediction algorithm, as in ref ^28^. Enzyme function can be grouped into different classes using the domain standard Enzyme Class (EC) ontology, which classifies enzymes into four levels based on the reactions they catalyse. The first level represents the general type of reaction (e.g., oxidoreductases), while subsequent levels add specificity, capturing information about substrates, cofactors, and products ^34^. This hierarchical structure provides as basis for relationships between enzyme functions and mechanisms that can be used to create biologically meaningful pairs.

To overcome the compute limitations with structural methods and existing benchmarks, we developed Squidly and a new benchmark for low sequence identity catalytic residue predictions. We combined a contrastive learning framework on PLM per-token embeddings with a rationally designed hierarchical pair scheme to create a sequence-based catalytic residue predictor that is accurate, scalable, and able to generalise in the absence of sequence similarity. Squidly exceeds the performance of both BLAST and SOTA ML tools in existing benchmarks achieving a F1 of greater than 0.85 across enzyme families, in addition to generalising to low sequence identity enzymes, with an F1 of 0.64 on sequences with less than 30% identity. Squidly is over 100x faster than folding a structure, enabling it to be used to annotate existing databases, in metagenomics pipelines, and to filter *de novo* designed proteins. Finally, owing to the added interpretability of sequence similarity-based approaches we provide Squidly as an ensemble with BLAST, and suggest using Squidly where the sequence identity is less than 30%. We envision Squidly will be broadly applicable in the enzyme engineering field and will complement existing sequence homology and structure-based methods.

## Methods

### AEGAN training and test datasets

To ensure comparability with the SOTA tools for catalytic residue prediction, the same training, test, and benchmark datasets (Uni14230, Uni3750) were utilised as described in prior work ^10^. These were originally sourced from UniProt and M-CSA databases. Uni14230 and Uni3750 datasets also include computational and manually curated predictions from SwissProt. To reduce redundancy, the data was filtered to retain only sequences with less than 60% sequence identity, resulting in 8,784 sequences in the training set and 1,955 sequences in the test set. Additionally, Squidly was evaluated against six previously reported benchmark datasets ^35–38^. The EF family, superfamily, fold, and HA superfamily datasets were specifically designed to include one representative sequence from each enzyme family, fold, or superfamily present in existing databases, and are thus not designed to predict “novel” active sites ^35,36^. The NN and PC datasets were both constructed to be structurally and functionally heterogeneous based on SCOP and CATH classifications ^37,38^. These datasets were generated prior to 2007 and are therefore limited in their catalytic diversity compared to the current databases. Both AEGAN and SCREEN were originally benchmarked against distinct subsets of these datasets, however, we found that they contained sequences with high identity to their respective training data, as detailed in Supplementary Table 2. Squidly’s reported performance with these datasets is based on the Uni14230 training set to ensure a fair comparison with AEGAN and given the lower overall sequence similarity between the training and test sets. An ablation of Squidly’s pair scheme was performed with single independent models using the Uni3175 test set. All other benchmarks report Squidly’s ensemble performance.

### New benchmark: CataloDB

To address the shortcomings of existing benchmark datasets for the evaluation of catalytic residue predictions, we introduce a benchmarking dataset named CataloDB. CataloDB collects the annotation data available for enzymes in the SwissProt database. The total set of enzymes includes experimentally determined annotations in addition to reviewed sequences with known structures, see Appendix A.1 for more details. Sequences with greater than 90% sequence similarity, based on shared overlap, were removed using mmSeqs2 ^39^. Structural and sequence-based clustering was performed on the filtered dataset to create a test set of 232 sequences with less than 30% sequence and structural identity to the training set (the remaining 5357 sequences). Structural identity was calculated using FoldSeek ^40^. Sequences that were in any of the six commonly used benchmarks were omitted from the training data to ensure compatibility between CataloDB and existing benchmarks. Additional detailed descriptions of the data are available in the Appendix A.2 and see Figures 3F and 3G for distribution of amino acids and EC classes in the benchmark dataset.

To enable direct comparison with the SOTA tool SCREEN, we retrained a lightweight version of the model using the CataloDB benchmark training data. This version of SCREEN, provided by the original authors, omits costly evolutionary features which are cumbersome for larger datasets. The authors report that these features have minimal impact on performance ^9^. Minor changes were made to the source code to improve model training stability. Notably, we allowed optimal stopping to occur after 5 epochs of training, rather than after 200 epochs in the original source code as we observed exploding gradients at later epochs when trained on CataloDB.

### ESM2 embeddings

Each sequence was encoded by either the 3 billion parameter or 15 billion ESM2 model ^23^. Pre-trained model weights were used without fine-tuning and per-token embeddings were extracted from the final layer of the encoder. These embeddings contain a vector of length 2560 (3B) or 5120 (15B) for each amino acid in the sequence. No normalization was applied to the embeddings before downstream use.

### Contrastive model

The supervised contrastive model is a network that takes per-residue embeddings as input (2560 or 5120 length vectors) and is trained to output a new embedding (N=128) of the residue. The model contains an initial dropout layer connected to the inputs, with a dropout rate of 10%, and two subsequent hidden layers of 1280 and 640 size each. During training, batches of 16,000 pairs are passed through the model and the distance between the output representations of each pair is calculated. The following cosine embedding loss function implemented by PyTorch is used to calculate the loss based on the distances:

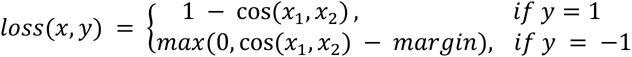

Where *x*_1_ and *x*_2_ are input vectors and *y* is the label, with *y* = 1 indicating a positive pair and *y* = *−*1 indicating a negative pair. When *y* = 1, the loss function rewards the model for maximising the cosine similarity, cos(*x*_1_*, x*_2_). The loss function minimises the cosine similarity when *y* = *−*1. Importantly, the pair selection scheme optimises for hard-negative pairs, and thus the margin is set equal to 0 when defining the loss function.

### Reaction informed pair-mining

Positive and negative pairs were created to enable contrastive comparisons where in the simplest form positives are pairs of exclusively catalytic or exclusively non-catalytic residues while negatives include a catalytic and a non-catalytic residue. As all possible pair combinations of amino acids become intractable, we are required to reduce this to make training feasible, hence we test three methods to subsample from this pool of possible pairs. Specifically, we used information about the EC number of the sequence and amino acid characters to refine the pair selection process. For further information, see Appendix A.2.

Three pair schemes were created to test for the effect of the pair scheme, herein referred to as “reaction-informed pair-mining”. See Figure 1.B for a visual description.

**Fig. 1:**
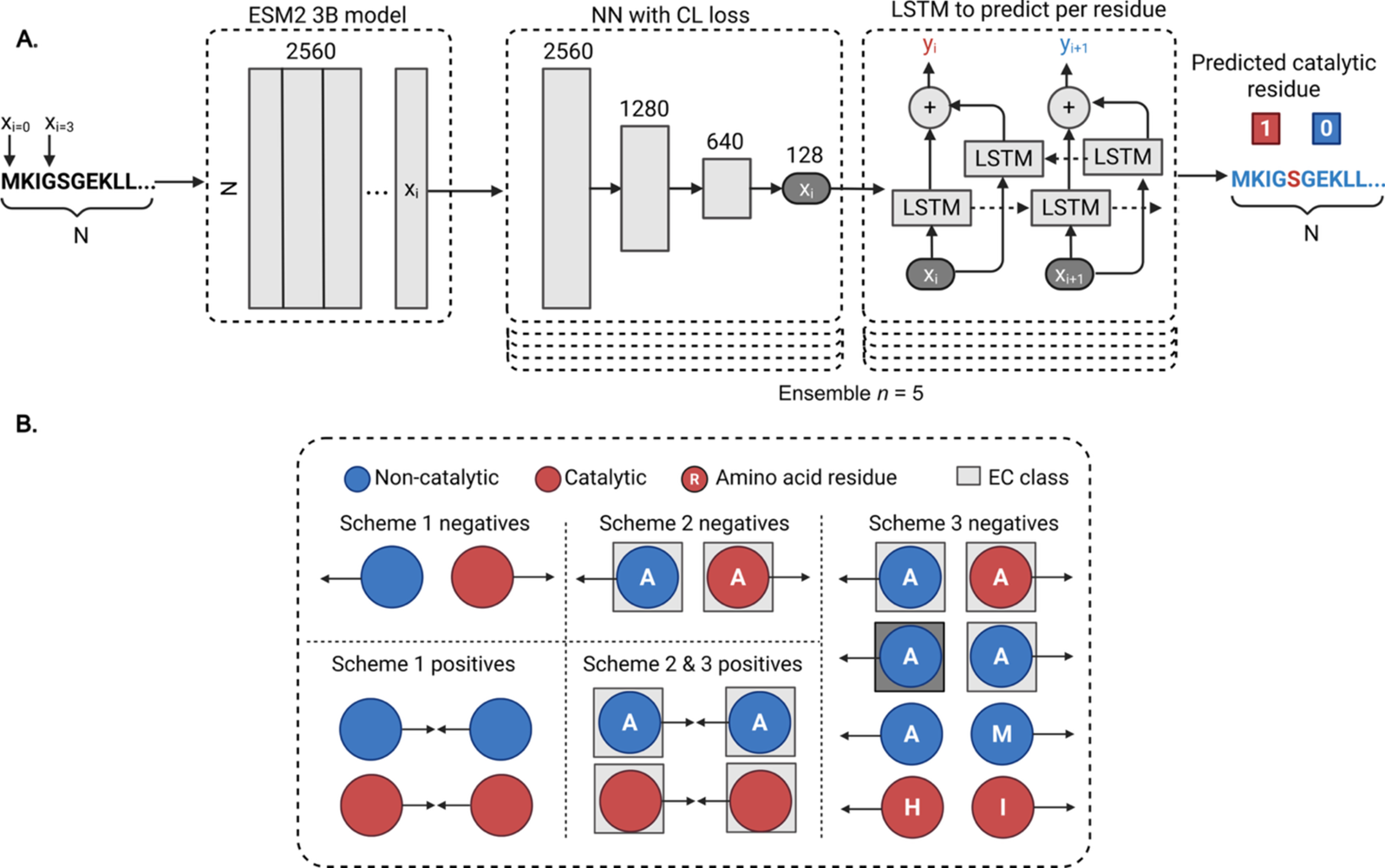
Overview of the model. A. Depicting a sequence being translated into per-token embeddings via ESM2, each per-token embedding is then translated into a smaller representation space by the MLP trained with contrastive loss (CL). Each residue from the CL model is then predicted as catalytic or not catalytic via a bidirectional LSTM network. The CL and LSTM models are ensembled (5 models) to improve performance and capture variance between the models. B. Representation of the pair-mining schemes depict the negative and positive classes for each scheme. Scheme 1 is the most permissive and is amino acid and EC *unaware*. Schemes 2 and 3 are more restrictive, requiring both the amino acids and the EC classes to be shared or different in the pair-mining scheme. Scheme 3 included additional negative pairs to separate EC numbers in the latent space.

Scheme 1 samples pairs across all sequences and residues such that positive pairs are pairs of residues that are either both catalytic or both non-catalytic.

Scheme 2 samples positive and negative pairs much like in scheme 1, except that each pair of residues must be the same amino acid and must have come from a sequence which has the same EC number.

Scheme 3 performs the same pair sampling as in scheme 2; however, scheme 3 also includes additional negative pairs that are either exclusively catalytic residues or exclusively non-catalytic and come from different EC numbers or amino acids.

To combat the inherent bias in the data towards certain EC numbers, sequences were grouped by EC number at the 2*^nd^* level, and an upper limit of 600 catalytic residues and 600 non-catalytic residues were sampled from each EC number. An equal number of positive and negative pairs were generated for each EC number. A parameter called the sample limit was set to limit the total number of pairs made for each group. The sample limit chosen was 16,000, providing a total of 8-10 million pairs for schemes 2 and 3. Scheme 1 was then trained with an equal total number of pairs.

### LSTM classifier

All LSTM classification models were trained using the same parameters. The models are bidirectional LSTM networks designed for binary classification of catalytic versus non-catalytic residues in protein sequences. The model takes as input 128-dimensional per-residue embeddings (from the contrastive model) that are processed by a 2-layer LSTM (hidden size: 128) with a dropout rate of 0.2, followed by a linear layer mapping the output to a single logit score, see Figure 1.A. Class imbalance is addressed by weighting the misclassification of catalytic residues 100 times higher during loss computation. The models were trained using a 90:10 train-validation split, with a batch size of 400 sequences, a learning rate of 0.01, and early stopping of 5 epochs was used. These parameters were not optimised. See Figure 1.A for a depiction of the model architecture.

### Squidly ensemble

For each training dataset, five models were ensembled by taking the per-residue output from the LSTM and then combining these to compute the mean score, along with the predictive entropy for each residue^41^. Users can specify a threshold for classifying a residue as catalytic (between 0 to 1.0): if the mean score exceeds this threshold, the residue is designated as “catalytic”. Users can optionally use the variance across the predictions to further filter for increased precision. The default thresholds were determined using the Uni3175 test datasets, see Supplemental Figure S3.

### BLAST similarity-based search

BLAST is a local alignment search tool able to identify homologous sequences from a reference database^42^. For our study, we utilised diamond BLAST to search the training set for similar sequences to each test set ^43^. The ultra-sensitive flag was used to identify sequences with low sequence similarities. Upon retrieval of results, the sequence with the highest sequence similarity was retained. This sequence was then aligned against the query using the alignment tool ClustalOmega ^44^. The gapped alignment was used to infer the catalytic residues of the query sequence.

## Results

### ESM2 per-token embeddings contain catalytic residue specific variation

To explore the information content contained within per-residue embeddings associated with catalytic residues, we performed principal component analysis (PCA) on equally sampled catalytic and non-catalytic residues from EC 3.1 and 2.7, Figure S1. For EC 3.1, the first two principal components explained 5.24% and 1.86% of the variance for all residues, respectively. Although the explained variance in PCs 1 and 2 is minimal, the separation in PCA space suggests that there may be some variation related to catalytic roles, Figure S1. The separability from a linear transformation suggests that ESM2 per-residue embeddings could be used to capture catalytic-specific signals.

### Reaction informed pair-mining improves upon contrastive learning and state of the art prediction performance

We hypothesised that contrastive learning would present an effective approach to leverage the rich feature space of ESM2 embeddings to distinguish between catalytic and non-catalytic roles. We implemented a biologically informed hierarchical contrastive learning pair selection framework to select pairs which are informative at the per-residue level.

### Ablation of pair selection as a key design choice in Uni3175 performance

To demonstrate the impact of pair-scheme design choices, we systematically evaluated the performance on the Uni3175 dataset of 1) a baseline approach using ESM2 embeddings, 2) the standard approach of randomly selecting pairs, versus 3) our proposed reaction-informed pair-mining schemes. Scheme 1, see Figure 1.B, is a “control” scheme and uses a standard contrastive loss function with random pair selection. We observed that scheme 1 performs poorly overall on the Uni3175 dataset, Figure 2.A. Across all datasets, Scheme 1 achieved relatively low F1 scores, with a mean F1 score of 0.49 and 0.43 for the 3B and 15B ESM2 models. Scheme 1 failed to surpass the LSTM classification model using unchanged ESM2 embeddings, which had a mean F1 score of 0.73, Figure 2.A. Schemes 2 and 3, both of which employ reaction-informed pair-mining, exhibited improved performance relative to both Scheme 1 and the ESM2 baseline. Scheme 2 achieved mean F1 scores of 0.80 and 0.82 using the 3B and 15B ESM2 models, respectively. Scheme 3 improves upon scheme 2 by contrasting catalytic and non-catalytic sites based on distinct EC and amino acid labels, Figure 1.B. This resulted in mean F1 scores of 0.79 and 0.84. The best performing model had an F1 score of 0.86 achieved using Scheme 3 and the 15B ESM2 model. Thus, scheme 3 is used for the ensemble model of Squidly and reported in all future benchmarks.

**Fig. 2:**
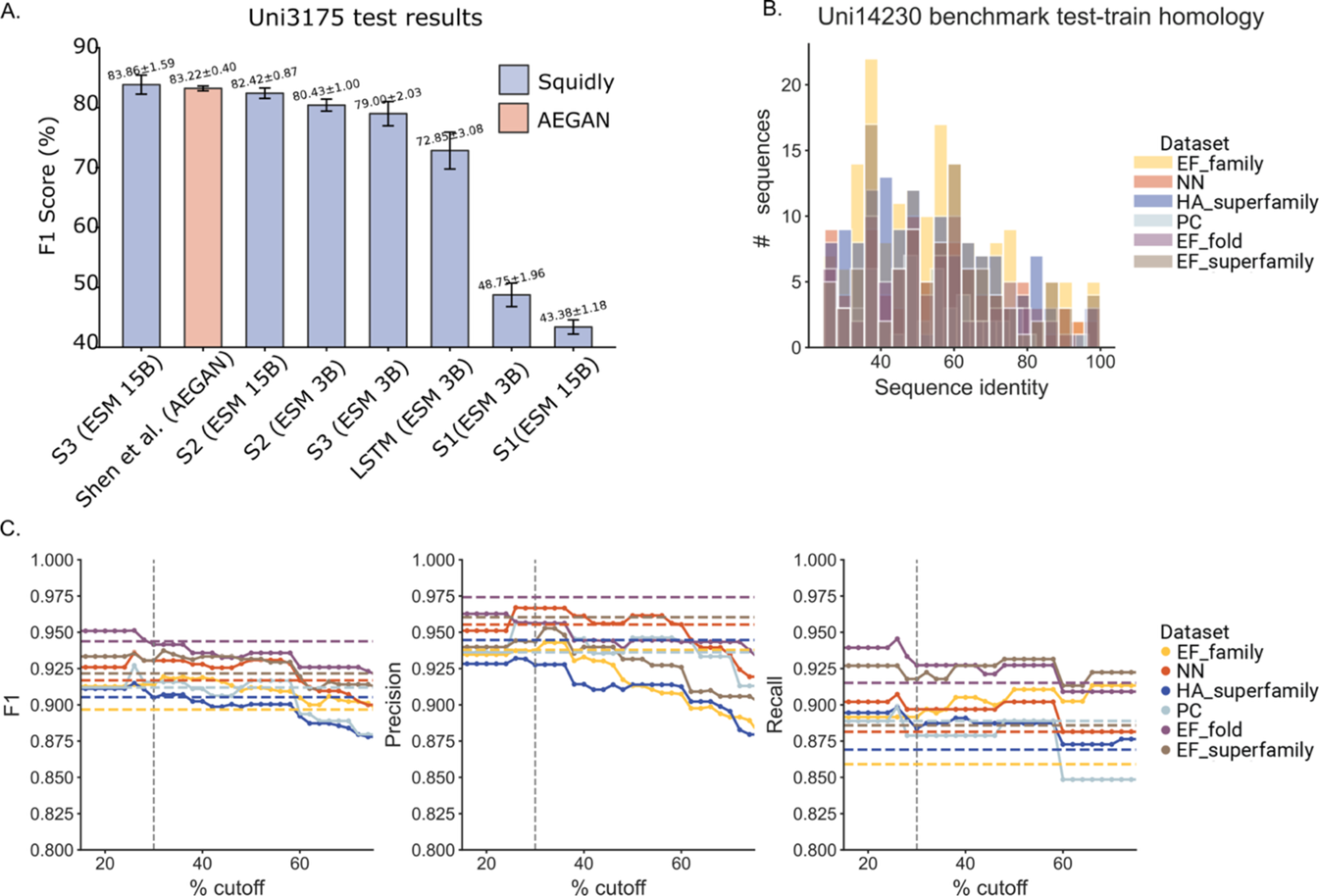
Performance of Squidly pair-mining schemes on the Uni3175 dataset. A. We compare the Squidly schemes without the ensemble with AEGAN, the tool by Shen et al., the authors of this dataset ^10^. Reaction informed schemes (Schemes 2 and 3) achieved the highest F1 scores, slightly surpassing AEGAN when using the 15B ESM2 model. The 3B ESM2 model remains competitive at a much lower computation cost. In comparison to the rationally informed pair schemes, the Scheme 1 (random pairing, S1) and raw embedding LSTM models performed poorly. B. Sequence similarity based on sequence identity to the closest sequence in the training set between each family. C. F1, recall, and precision for the ensemble of Squidly (3B, scheme 3) and BLAST with different sequence identity cut-offs. The cut-offs indicate the point at which to transition from a Squidly to a BLAST prediction. At 100% only Squidly is used for predictions, and at 0%, BLAST is used, unless there are no similar sequences. For sequences with less than 30% sequence identity (dashed black line), Squidly predictions are used. The dashed horizontal line represents the total score for BLAST on the specific dataset, while the dots represent the combined BLAST and Squidly ensemble approach. Squidly reports higher scores for recall, while BLAST is more precise. The performance is best when used as an ensemble.

The results from Uni3175 benchmark show that Squidly improved marginally upon the performance of previous work and the current SOTA ML prediction model AEGAN by Shen et al. ^10^ Squidly uses only sequence through ESM2 embeddings for prediction, which, compared to the homology and structural modalities included in AEGAN, makes for a more efficient predictor without compromising performance.

We additionally benchmarked Squidly with the multiclass classification benchmark from EasIFA on a further 66 sequences with less than 40% sequence identity to either training set ^11^, see Appendix 4 for more details. EasIFA performs best, with a recall of 79.05% and precision of 90.22%. Squidly had a recall of 78.10% and precision of 73.87%.

Overall, the results on the Uni3175 dataset indicate that the rationally informed pair-mining strategy is an effective method for discriminative tasks using per-token ESM2 embeddings. It also indicates that sequence-based models using per-token PLM embeddings can compete with models using structural modalities. See Methods for details on each scheme design.

### Squidly outperforms other ML-based methods yet works best in conjunction with sequence similarity methods

Squidly was evaluated against six previously reported benchmark datasets, namely the EF fold, family, and superfamily benchmarks, as well as the HA superfamily, NN, and PC benchmarks, see Supplementary Table 1 and Appendix 4 for description of each dataset, see Figure 2.B for sequence identity between test and train sets. Squidly’s performance exceeded those of SOTA models AEGAN and SCREEN, Table 1. When compared to the baseline of BLAST, we find that BLAST outperforms all other ML models except Squidly, likely due to the homologous sequences in the test and train sets, see Appendix A3, and Supplementary Table 2 for details.

**Table 1.** F1 scores of tools on 6 common benchmark datasets.

| Tool | EF-fold | EF-superfamily | EF-family | HA-superfamily | NN | PC |
| --- | --- | --- | --- | --- | --- | --- |
| PREvail<sup>17</sup> | 0.2579 | 0.2575 | 0.2585 | 0.2483 | 0.2537 | 0.2582 |
| CRpred<sup>8</sup> | 0.2613 | 0.2643 | 0.2651 | 0.2628 | 0.2638 | 0.2630 |
| AEGAN*<sup>10</sup> | 0.8369 | 0.8687 | 0.8552 | 0.8259 | 0.8177 | 0.8736 |
| SCREEN*<sup>9</sup> | 0.645 | 0.615 | - | 0.72 | 0.738 | 0.741 |
| Squidly (ESM2 3B, S3) | 0.8631 | 0.8775 | 0.8545 | 0.8171 | 0.8354 | 0.8527 |
| Squidly (ESM2 15B, S3) | 0.8765 | 0.8936 | 0.8867 | 0.8513 | 0.8586 | 0.8454 |
| BLAST (ultra-sensitive) | 0.9437 | 0.9216 | 0.8967 | 0.9053 | 0.9169 | 0.9119 |
| BLAST + - Squidly (ESM2 3B, S3) | **0.9509** | **0.9333** | **0.9126** | 0.9111 | **0.9259** | **0.9119** |
| BLAST + - Squidly (ESM2 15B, S3) | 0.9476 | 0.9274 | 0.9034 | **0.9179** | 0.9230 | 0.9119 |

Benchmark results for PREvail, CRpred and AEGAN were sourced from <sup>10</sup>. Results from Screen were sourced from <sup>9</sup>. BLAST results were performed using the ultra-sensitive flag and aligned to the closest recorded enzyme with an active site in the training dataset (Uni14230). Squidly results shown for models trained on the Uni14230 dataset (same as AEGAN) using pair mining scheme 3 (S3). \* Indicates structure-based tools.

We hypothesised that BLAST would work best in instances where a similar sequence existed while Squidly would excel in low sequence similarity situations. To test for the optimal conditions for integration we varied the rate at which either predictor was used, Figure 2.C. For sequences with low sequence identity (<50%), using Squidly improves the F1 score. However, Squidly has a higher likelihood than BLAST to record false positives where similar sequences exist while BLAST is more precise. The cutoff of 50% sequence identity was used to compare the performance of the ensemble method to existing tools. On all datasets using this cutoff an ensemble of BLAST and Squidly performs SOTA, Table 1.

### Squidly is scalable and fast

Squidly’s sequence-based approach reduces the overall computational cost of prediction to enable the screening of large databases. SOTA ML models rely on accurate structures as input, and thus structural prediction must be done prior to inference when screening databases such as metagenomic samples. Squidly’s smaller 3B model can be run locally and can predict roughly 10 sequences a second in our tests using an NVIDIA H100 GPU. In comparison to the current widely used structural generation tool Chai ^45^, Squidly is approximately 200-fold faster (e.g. N=100, Squidly=44s, Chai=13263s). Note this under- represents the time for the structural tools to run, as they also require further prediction which also adds time, Table 2.

**Table 2.** Resource usage measurements for Squidly models compared to structure prediction tool Chai-1 when predicting 100 sequences with an average length of 443 residues.

| Model | Wall Clock (s) | VRAM (GB) | RAM (GB) |
| --- | --- | --- | --- |
| Squidly 3B | 43.9 | 26.5 | 17.1 |
| Squidly 15B | 185.6 | 74.8 | 91.5 |
| Chai-1 | 13263.0 | 20.8 | 19.3 |

All models benchmarked using the Nvidia H100 GPU with 80Gb of VRAM, AMD EPYC 7643 48-Core Processor on a HPC cluster running Rocky Linux 8.10.
Squidly models are ensembled ( $n = 5$ ).

### Squidly generalises to low identity sequences

To evaluate the capacity of Squidly to generalise to low identity sequences, we developed a benchmark dataset, CataloDB with low structural and sequence identity, see Methods and Figure 3.A, B for similarities between the test and train sets. This benchmark serves as an alternative test set that takes structural and sequence similarity into account, allowing for fair comparison between future structural and sequence-based tools and testing in low identity settings. Squidly 3B and 15B models showed similar performance on the benchmark with most sequences (N=148/232, N=138/232), recording at least one correct catalytic residue. For comparison, BLAST identifies only 68 sequences that have a catalytic residue recovered correctly. Both Squidly models performed similarly with F1 scores of (0.66, 0.69), precision of (0.86, 0.81), and recall of (0.52, 0.61) for the 15B and 3B models (respectively), while BLAST records an F1 of 0.37, recall of 0.25, and precision of 0.69, see Figure 3.C. We also retrain SCREEN, with performances recorded above BLAST however still below Squidly, Figure 3.C, see Methods for details.

**Fig. 3:**
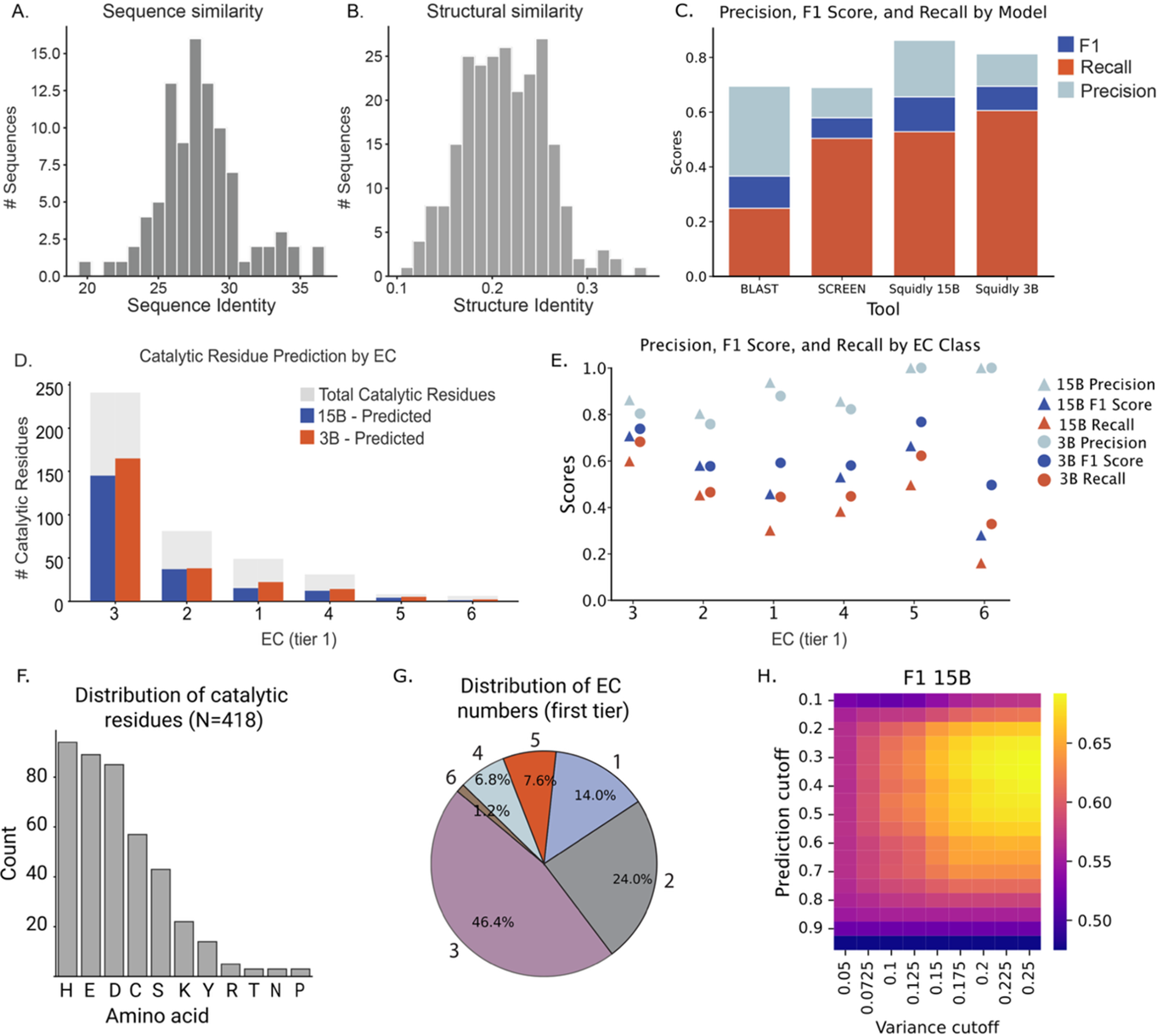
A. BLAST sequence similarity to the training set for low identity CataloDB test set sequences. B. Structural similarity for the final test set, filtered on both structure and sequence. C. F1, recall, and precision for Squidly (scheme 3), SCREEN and BLAST on CataloDB. D. The number of CataloDB catalytic residues predicted (true positives) by Squidly (scheme 3) for the 6 available EC classes relative to the decreasing total number of true catalytic residues. E. The Precision, F1 and Recall score of Squidly predictions for the 6 available EC classes. F. The distribution of the identity of the catalytic residues is not even and represents the uneven distribution of the catalytic mechanism within the benchmark dataset. G. Almost 50% of the data points are in the well characterised class of EC 3, which are hydrolases. Neither EC 7 nor EC 6 is represented, which are translocases and ligases, respectively. H. Performance of the Squidly 15B ensemble model on the CataloDB benchmark under varying prediction and variance thresholds. Note, default 0.5 prediction cutoffs were used to evaluate Squidly’s performance in C.

Reflecting the bias of public data, CataloDB shows an over-representation of certain EC numbers and amino acids. To examine the effect of this bias, we stratified Squidly’s performance across the EC classes and observed a measurable drop for less well-represented sequences. Despite this, F1 scores remained relatively stable for two of the underrepresented classes, ECs 5 and 6, see Figure 3.E. However, we note that EC 6 was represented by only three sequences in the test data. Across all ECs, the models displayed a consistent trend of higher precision than recall, see Figure 3.D. Overall, these results suggest that Squidly can generalise to low identity settings and be used in predictions where traditional homology transfer is not possible. However, caution should be taken when applied to poorly represented enzyme classes such as EC 6 and 7.

## Discussion

Rationally constraining pair generation in contrastive learning can improve model performance on biological data. Using publicly available EC number annotations and amino acid sequences, we categorised pairs of catalytic residues into distinct groups enriched with hard negatives to enable sequence-only prediction of catalytic residues. Squidly’s pair selection procedure produces distinct samples of the training data for training each model. The lightweight architecture of the per-token contrastive learning model, coupled with these independent data subsets, make Squidly well-suited to ensemble learning, as the diversity between model training data increases ensemble robustness and reduces correlated errors across models^41,46^. This approach led to substantial improvements compared to a standard LSTM classifier and typical pair-mining strategies. Squidly significantly outperformed other ML-based sequence methods and showed marginal improvement to SOTA ML-based catalytic residue prediction models that use structural data while running in a fraction of the time. Furthermore, Squidly performs comparably to BLAST when homologous sequences exist, yet also shows good performance in low-identity settings, where BLAST is unable to identify any similar sequences. Given the complementarity of BLAST and Squidly, we provide Squidly as an ensemble with sequence similarity, leveraging the interpretability of sequence similarity-based methods where homologous sequences exist, and Squidly, in out-of-distribution settings. The focus of this work was catalytic site prediction; however, of similar utility are combined binding and catalytic prediction models, such as EasIFA ^11^, and therefore future work should additionally consider binding sites.

A limitation of this study is the inherent bias in publicly available data towards well studied enzyme classes. The bias likely confounds the perceived performance on less studied mechanisms, and in these instances the results should be considered with caution. For example, when ensembling models, we observed that the optimal classification threshold varied across EC numbers and ESM model variants, indicating sensitivity to the underlying distribution. This suggests that out-of-distribution datasets explicitly designed to reflect the intended application domain are needed for fair threshold calibration. A further challenge is the lack of a consolidated benchmark for catalytic residue prediction, making direct and fair comparison between tools difficult. EasIFA showed superior performance on the overlapping test set (n=66), indicating when structures are available, EasIFA is a good tool for catalytic residue prediction. However, we note that the multiclass classification objective and benchmarks used to evaluate EasIFA ^11^, differences in sequence similarity to the test set, and small test set size limit these conclusions. Existing benchmark datasets do not capture the diversity of enzyme catalysis. Although we have provided a new benchmark that improves and expands upon previous datasets by using structural identity filtering, the core issues remain. CataloDB and current benchmarks are small and are likely enriched for model organisms with well-studied enzyme mechanisms. While low sequence and structural similarity samples are held out from training, they may not accurately represent the generalisability of these methods in out-of-distribution settings. Furthermore, the datasets often consist of sequences with known structures in the PDB, many of which were likely part of the AlphaFold training set or structurally similar clusters represented in the set. Tools also evaluate their performance using experimental, rather than predicted structures ^10^. Consequently, SOTA tools that rely on AlphaFold predicted structures tend to perform better in these benchmarks but are likely to underperform in practical applications involving truly unique and uncharacterised sequences. This issue also applies, albeit to a lesser extent, to the method proposed in this study, which utilises ESM2 representations. While ESM2’s training set is more diverse and comprehensive, it may not generalise to distant out-of-distribution sequences compared to the well-studied sequences used in the benchmarks. Although Squidly facilitates annotation in cases where similarity methods fail, its usability in practice should be considered provisional, as generalisability to less-studied enzyme lineages and catalytic mechanisms remain uncertain and may not extend beyond the well-represented mechanisms captured in current benchmarks.

Beyond these benchmark constraints, Squidly and related methods are constrained by the experimental annotations that underpin the training data. Currently there exists a limited set of experimentally validated annotations ^13^. Datasets such as Uni14230 and CataloDB used for training in this study include SwissProt sequences with reviewed computational annotations to increase the size of the training data. However, this does not improve the diversity of catalytic mechanisms and enzyme sequences, as these methods reflect the inherent biases in databases towards well represented EC classes. Squidly, having learnt from this biased data, may indeed replicate this bias during inference and is therefore less likely to provide insights into novel or understudied catalytic mechanisms. Despite the limited data for catalytic residue prediction, we show that an ensemble approach of machine learning and sequence-similarity-based methods performs SOTA across multiple benchmark sets. By leveraging contrastive learning, we find that ML sequence-based predictions can generalize to low sequence/structural similarity settings. Finally, the methods reported in this paper that leverage rational data engineering for ensemble contrastive learning have broad implications for learning across biological data. Biological sequences are generated through evolutionary processes occurring over billions of years, resulting in hierarchically structured data that has significant variance between evolutionarily distant sequences. Moreover, mechanisms in biochemistry are often similar for related chemical reactions. By leveraging ontologies such as EC numbers, pairs can be selected rationally to improve training on proteins that are not only evolutionarily related but also share broader biochemical functions. A wide range of alternative ontologies exist that can capture different structures within biological data, suggesting that this approach is likely to be applicable across other biological classification tasks, particularly in studies utilising contrastive frameworks for per-token in- sequence classification ^47^. Future studies could explore alternative pair combinations beyond EC numbers, as well as additional ontologies to better understand the utility of such data structures.

## Code and data availability

All source code is available at Github (https://github.com/WRiegs/Squidly). All associated data including CataloDB is available at Zenodo (https://doi.org/10.5281/zenodo.15541320).

## Author contributions statement

W.R. and A.M. conceived the experiment(s), W.R. designed and implemented the model, W.R. and A.M. ran the experiment(s), analysed the results, and wrote the manuscript. M.B. and F.H.A reviewed the manuscript and provided input to the research.

## Acknowledgments

A.M. is supported by the Schmidt Science Fellows, in partnership with the Rhodes Trust. W.R. is supported by the Australian Government Research Training Program (RTP) Scholarship. The authors thank Charlie Trimble for his generous contribution to the project.

## APPENDIX

### A.1 ESM2 per-token embeddings contain catalytic residue specific variation

To illustrate the information content of per-residue embeddings for catalytic residue differentiation, we performed principal component analysis (PCA) on equally sampled catalytic and non-catalytic residues from EC 3.1 and 2.7, Figure S1. For EC 3.1, the first two principal components explained 5.24% and 1.86% of the variance for all residues, respectively. Although the explained variance in PCs 1 and 2 is minimal, the separation in PCA space suggests that the largest two signatures of linear variation could be related to catalytic roles, catalytic (red-coloured) and non-catalytic (blue-coloured) residues, Figure S1. This exploratory analysis motivated us to test this hypothesis and proceed with per-residue embeddings for our contrastive learning model.

### A.2 Rationale for developing CataloDB

To enable comparisons to existing work, we utilized the AEGAN training and test datasets as described above. However, the limitations identified motivated us to create a new benchmark, CataloDB, for future evaluations. The pre-existing benchmarks were published prior to 2007 and were developed by selecting proteins with active site annotations representing unique protein families, superfamilies, or folds. Their construction lacked standardized metrics for ensuring appropriate similarity separation between training and test data. The continued use of these historical benchmarks likely persists to maintain backward compatibility in comparisons with earlier methods. However, these six benchmarks predominantly represent well-studied folds and families, making it challenging to create properly separated training and test sets.

Our results demonstrate that BLAST already performs comparably or better when predicting catalytic residues in sequences with greater than 0.35 identity. Therefore, we believe machine learning methods should aim to develop tools that generalize effectively to low-identity sequences. CataloDB addresses this challenge, and the concerns by containing test sequences with strictly less than 0.3 sequence and structure identity to the training set. This benchmark is more extensive than previous collections while still allowing for a larger training set with a higher redundancy threshold (90%), enabling machine learning methods to better learn from the underlying data distribution.

#### Evaluation of Swissprot data for use in CataloDB

To overcome the limited availability of experimentally validated catalytic residue annotations, annotations from sequences which have been reviewed by Swissprot’s expert review panel have been included in training and test data in recent machine learning tools ^9,10^. However, the assumption that these annotations are valid is not a given. Swissprot provides limited information to clarify how their review process for catalytic mechanisms is conducted ^12^. These annotations may have been generated manually or by computational predictions and likely contain the same bias present in the original experimentally validated set. Here we present initial analysis done, using an earlier version of Squidly, to determine whether reviewed sequences improved performance, and thus, the quality of training data.

Three datasets were curated to evaluate the impact of reviewed sequences on training catalytic residue prediction models. Dataset 1 contained 2,209 sequences with only experimentally validated catalytic residue annotations. Dataset 2 included 6,437 sequences, including experimentally validated annotations and sequences with catalytic residues supported by known structures. Dataset 3 included 99,926 sequences comprising all catalytic residue annotations in the SwissProt database. Redundancy reduction removed sequences with over 90% sequence identity. After filtering, Dataset 1 was reduced to 2,030 sequences, Dataset 2 to 5,921 sequences, and Dataset 3 had the highest proportion of redundant sequences with only 57,698 sequences left after filtering. See Table 2 for details.

Contrary to the expectation that increasing data size would improve model performance, we found that increasing the available training data by including reviewed sequences from SwissProt does not improve the overall bias or variance of the available experimental data. Figures S6, S8 and S10 show the EC distributions (EC numbers up to the second tier) across the datasets. All three datasets are biased toward hydrolases (EC 3), followed by classes 2, 1, 4, 5, 6, and 7. The relative abundance of EC class 2 increases between dataset 1, dataset 2, and dataset 3, while other classes remain relatively constant. This trend suggests that the additional sequences in the reviewed datasets likely originate from similarity-based prediction methods that incorporate experimental data, thus maintaining the relative abundance in EC representation despite increasing dataset size.

Figures S4, S6 and S8 compare the distributions of amino acids acting as catalytic residues across the datasets. The distributions are dominated by histidine, aspartic acid, glutamic acid, cysteine, and serine, in line with the dominant classes being hydrolases. Dataset 2 and dataset 3 show a relative increase in aspartic acid abundance compared to dataset 1. Despite this, and the much larger size of datasets 2 and 3, their amino acid distributions closely resemble dataset 1, indicating that reviewed datasets contribute little diversity in rare catalytic residues, which may correspond to mechanisms that are less studied. Overall, these findings suggest that training the models using the additional data will not greatly improve the performance of the models when generalising to sequences with low identity in the experimental ground truth set. Based on our analyses we use dataset 2 for CataloDB.

### A.3 Reaction-informed pair schemes

By performing contrastive learning with pairs, we move from having too few samples to having too many samples. The large pool of amino acid pairs in our dataset varies in how informative each pair is for the model. Therefore, we must find the most effective ways to down sample and select the most informative pairs for our model to learn from. Particularly, we want to maximise the proportion of hard negative pairs in the training data.

The Uni14320 training dataset contains 8,735 non-redundant sequences from Dataset 1 (experimentally validated) with an average sequence length of 407 residues. Given the total pool of 3,555,145 residues, a total number of 6.32 × 10^12^ unique pairwise comparisons can be made, as determined by the binomial coefficient 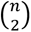. We developed pair schemes 2 and 3 to capture the most informative pairs from this available data. Maximising hard negatives improves the model’s ability to discriminate catalytic and non-catalytic residues in a more generalisable way. If two pairs of residue embeddings are inherently very distinct from one another, the model will be unlikely to learn much from them. This is not just because our loss function will penalise the model less during training, but because the relationships extracted from the feature space that are used to separate these two inputs might not necessarily be related to our primary objective.

Reaction informed pair-mining assumes that there must be key biological contexts which create harder negative pairs for the model to differentiate. First, we make the basic assumption that the model will be unable to correctly discriminate between similar amino acids pairs. It is likely that a contrastive model will find pairs individual amino acids, such as histidine, harder to contrast, because the unique structure and properties of each amino acid determine the catalytic or non-catalytic roles in which these amino acids play within enzymes. Therefore, we decided to implicitly include the amino acid character of each catalytic site into the pair sampling scheme.

Another assumption we made was inspired by the use of contrastive learning in another catalytic site prediction tool developed by Tong Pan et al. ^9^. Their contrastive model creates EC specific representations of sequences for further use in a convolutional graph neural network classification model. Their contrastive model, however, did not try to use these contexts to better differentiate between catalytic and non-catalytic sites in their model, but as a support to ensure the information about the sequence EC is propagated through their multimodal system. EC numbers separate enzymes based on the reactions they perform. Not only that, but lower-level EC numbers group enzymes by the types of substrate bonds they cleave, or mechanisms by which they catalyse reactions. Inspired by this idea, we leveraged EC numbers to sample hard negative pairs. We specifically chose to represent sequences with only the first two levels of their EC-numbers so that there is sufficient variety in the data that represents each label.

### A.4 Evaluation of Existing Benchmarks

The six standard benchmarks traditionally used for catalytic residue prediction exhibit methodological inconsistencies in the literature. In our comparative analysis, we discovered that state-of-the-art methods benchmarked in this study – specifically SCREEN and AEGAN utilized different independent subsets of these benchmarks, complicating direct performance comparisons. According to the SCREEN authors, data preparation involved clustering the EF family and M-CSA training data to less than 0.4 sequence identity, selecting unique cluster representatives for training and validation. The authors reported that they “excluded enzymes from these five test datasets from our training/validation dataset.”

Our analysis identified that up to 85.4% of the test sequences had greater than 0.9 identity to the training set in the already filtered subsets provided by the authors. Notably, the high sequence similarity appears reduced in the EF superfamily and EF fold datasets, likely resulting from the authors’ use of the closely related EF family dataset during training set curation. The test results published by SCREEN align with our observations, showing a marked increase in F1 score when comparing the HA, NN, and PC datasets with the EF datasets, corresponding to their differing levels of data leakage.

Additionally, the curation process employed in SCREEN for the training data appears unnecessarily stringent, with the average closest training sequence similarity measured at only 0.21. This sparse training data likely impacts machine learning models attempting to fit such data distributions effectively. We hypothesize this factor may explain why AEGAN reportedly performed poorly when retrained using this data in the SCREEN authors’ experiments.

### A.5 Benchmarking Squidly against EasIFA multiclass classification

EasIFA provides both a multiclass classification model and a binary classifier that labels residues as active sites. The binary labels include catalytic, binding, and “other” residues from UniProt, whereas Squidly, AEGAN and SCREEN are trained specifically to identify catalytic residues. This definitional difference makes direct comparisons challenging, owing to the difference in training dataset between the models.

Here we re-evaluated the performance of EasIFA, AEGAN, and Squidly using the pretrained EasIFA multiclass classifier and its benchmark dataset. To minimize overlap with training data, we retained only sequences that had <40% sequence identity to the Squidly and AEGAN training data (Uni14230) and to EasIFA’s training data, and that contained at least one annotated catalytic residue. After filtering, 66 sequences remained, these include no representative sequences from EC classes 4, 6 and 7, See Supplemental Figure 2.B. The subset had an average identity of 0.082 to the Squidly/AEGAN training sets and 0.238 to EasIFA’s training set.

In the EasIFA manuscript, recall was calculated as zero for sequences without catalytic residues, which artificially reduced the reported averages. As such we recalculated recall per-sequence after removing sequences with no catalytic residues.

The benchmark yielded the following outcomes. EasIFA achieved a recall of 79.42 and a precision of 79.55, while the best-performing Squidly model reached a recall of 77.40 with a precision of 65.76. AEGAN recorded a high recall of 91.78 but an unusually low precision of 7.85, which may indicate a benchmarking artefact rather than an inherent property of the model. Across the nine independently initialized Squidly models tested, performance showed considerable variance, with recall averaging 58.74% (standard deviation 11.06%) and precision averaging 54.00% (standard deviation 6.82%). EasIFA’s benchmark did not provide variance estimates, suggesting that the best-performing models were also reported there.

Squidly and AEGAN tend to agree with each other, having on average the same true positive and false positive predictions 96.20% of the time (TP=100.00%, FP=92.41%). Many of the false positives predicted by Squidly were binding sites, See Supplemental Figure 2.C. This tendency is perhaps because of the overlap between binding and catalytic residue annotations sometimes found in UniProt. Overall, EasIFA shows strong performance in this benchmark. However, due to the dataset limitations, differences in training data and label definition, and EasIFA’s higher sequence identity to the test set, these results should be interpreted cautiously. This benchmark shows that EasIFA may have improved precision compared to Squidly and AEGAN. Additionally, the EasIFA model can predict binding sites alongside Catalytic Residues, which is advantageous when studying cofactor-dependent enzymes using EasIFA’s method.

---

**Fig. S1:**
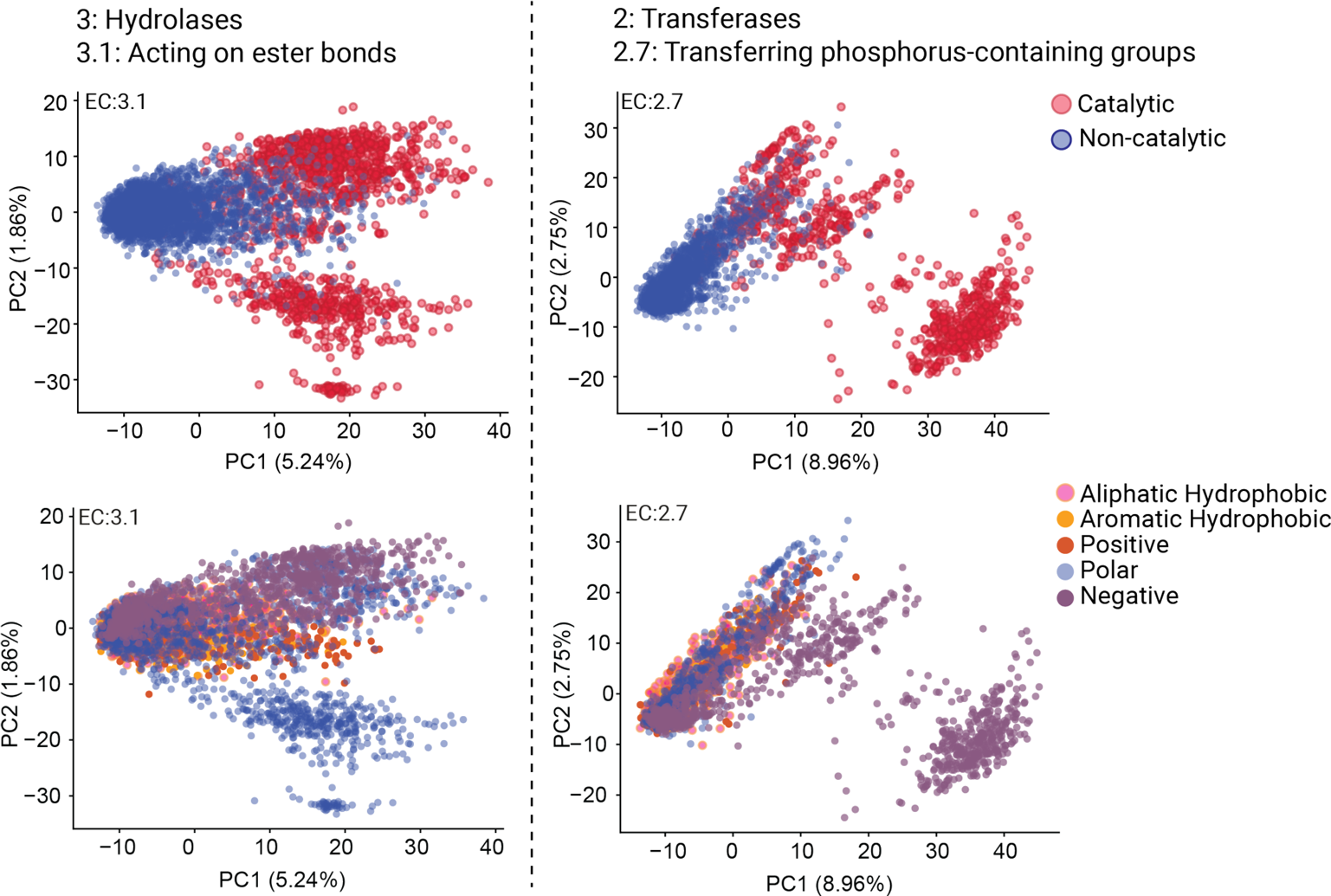
Principal component analysis of per-residue embeddings from EC numbers 3.1 and 2.7. In the upper row, red colours indicate a catalytic residue, and blue, non-catalytic. In the lower row, residues are coloured by amino acid class which represent their structural and chemical properties. The x and y axes represent the first and second principal components in each subplot. A clear separation of catalytic and non-catalytic residues is seen in the principal components which explain the most variance in our data.

**Figure S2:**
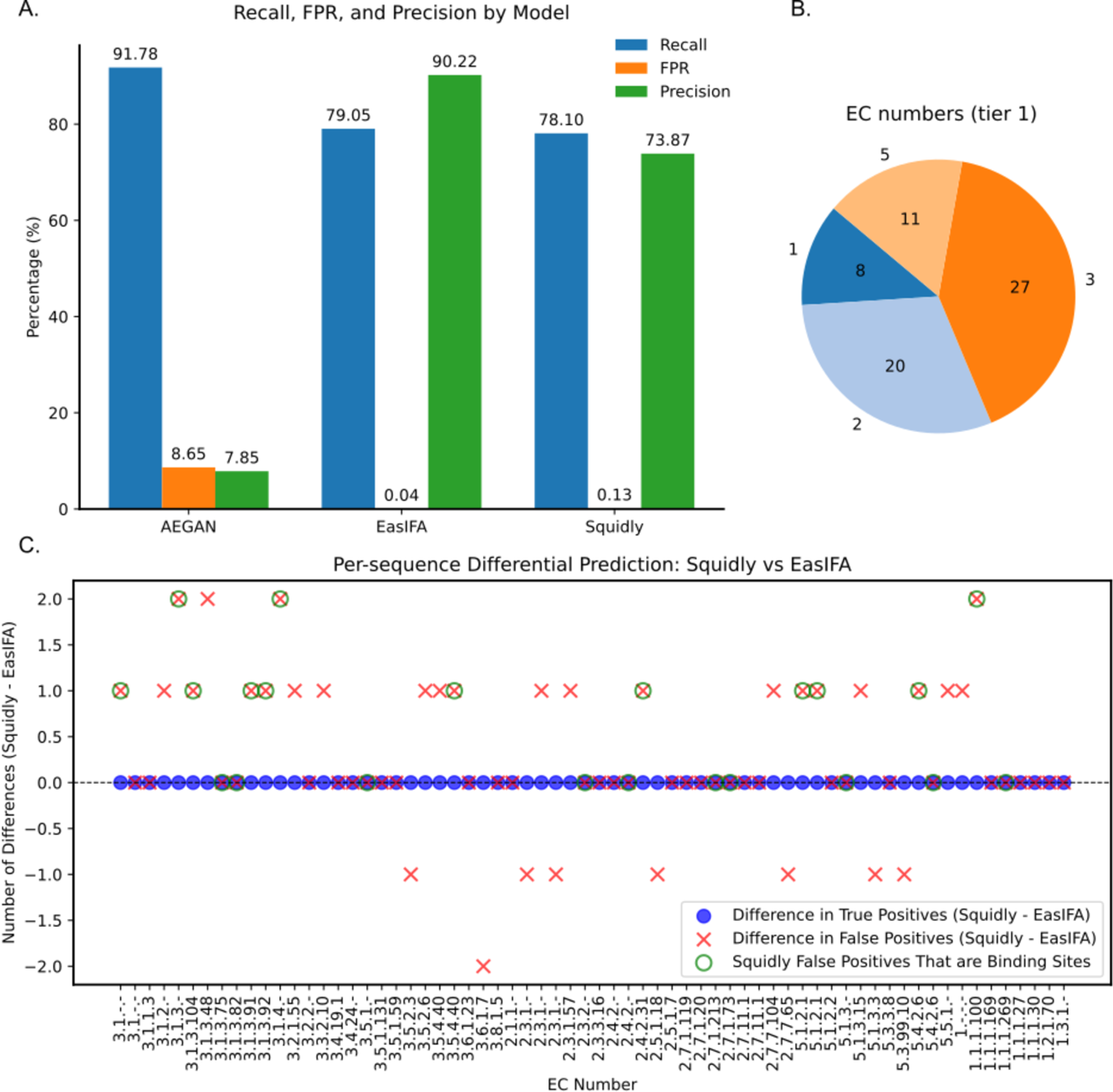
A. Recall, precision, and false positive rate (FPR) for AEGAN, EasIFA, and Squidly on the filtered EasIFA test subset. EasIFA shows balanced performance, Squidly is competitive given its sequence-only inputs, and AEGAN displays unusually low precision that may reflect a benchmarking artefact. B. Most of the test set are sequences from EC classes 2 and 3, with no representation for classes 4, 6 and 7. C. The differences in false positives, and true positives predicted by Squidly’s best model and EasIFA. Although the models agree with 100% of true positive predictions made by both tools, Squidly has a higher tendency for false positives, with many of the false positives being true binding sites.

**Figure S3:**
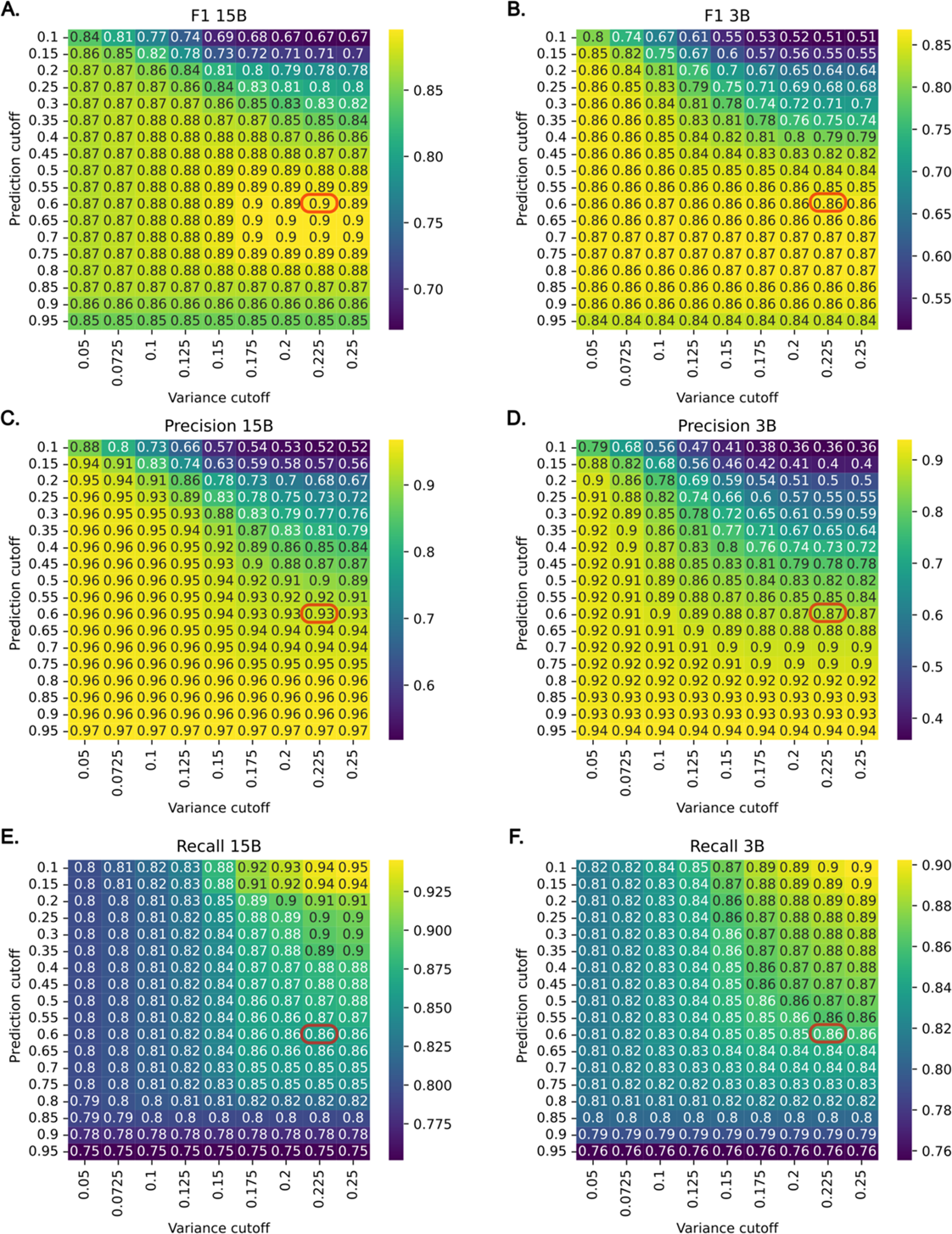
The default ensemble threshold for prediction and variance cutoffs of 0.6 and 0.225 (outlined in red) were selected to balance the precision and recall of the Squidly ensemble model on the Uni3175 benchmark under varying mean prediction and variance thresholds. Shown are F1 scores, precision, and recall for the 3B and 15B models. A. F1 of the 3B model, peaking at 0.70. B. F1 of the 15B model, peaking at 0.69. C. Precision of the 3B model shows that as the cutoff increases. D. Precision of the 15B model. E. Recall of the 3B model. F. Recall of the 15B model.

**Figure S4:**
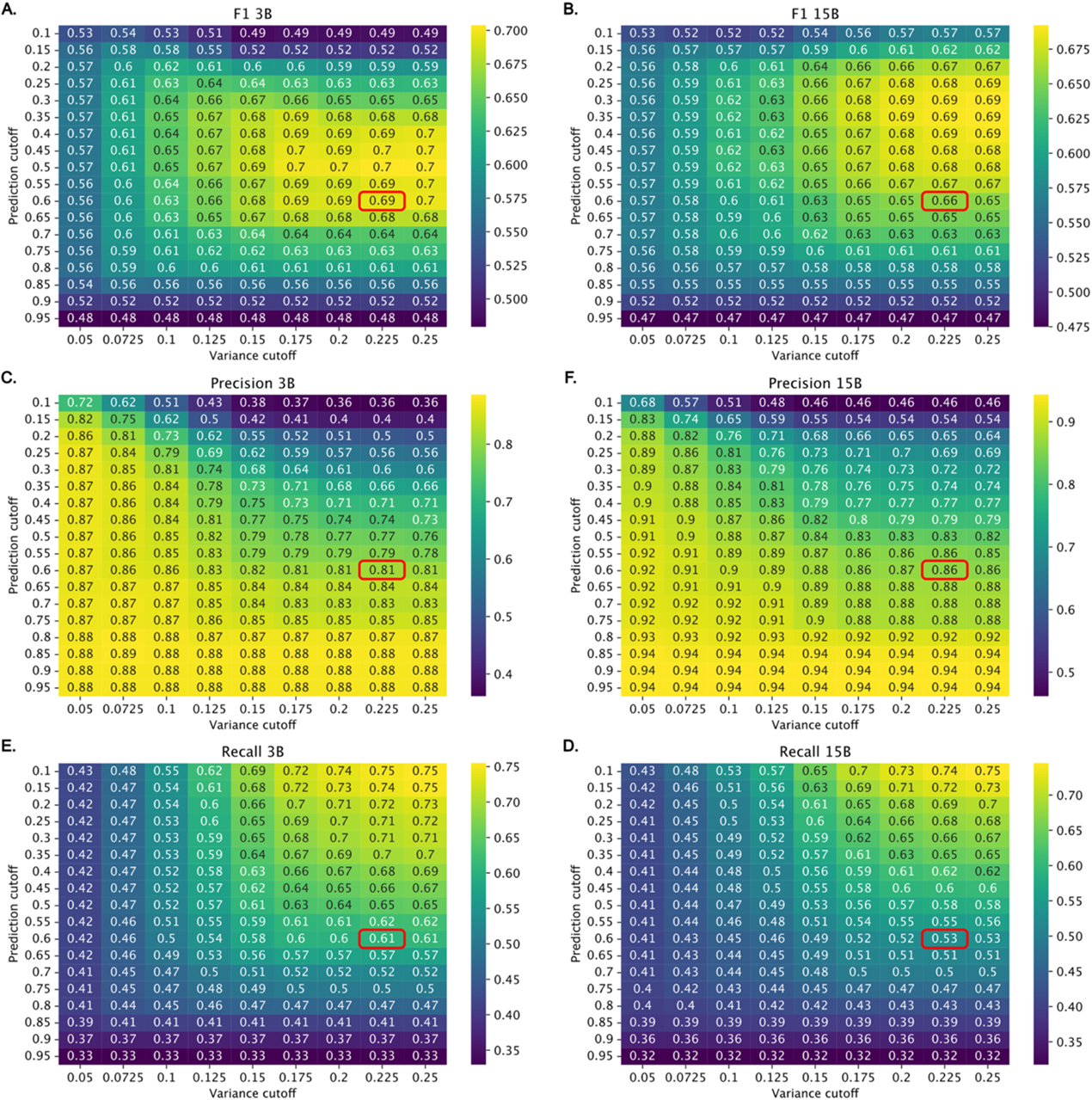
Performance of the Squidly ensemble model on the CataloDB benchmark under varying mean prediction and variance thresholds. Shown are F1 scores, precision, and recall for the 3B and 15B models. A. F1 of the 3B model, peaking at 0.70. B. F1 of the 15B model, peaking at 0.69. C. Precision of the 3B model shows that as the cutoff increases. D. Precision of the 15B model. E. Recall of the 3B model. F. Recall of the 15B model. The default ensemble threshold for prediction and variance cutoffs of 0.6 and 0.225 (outlined in red) were selected using the Uni3175 benchmark.

**Figure S5:**
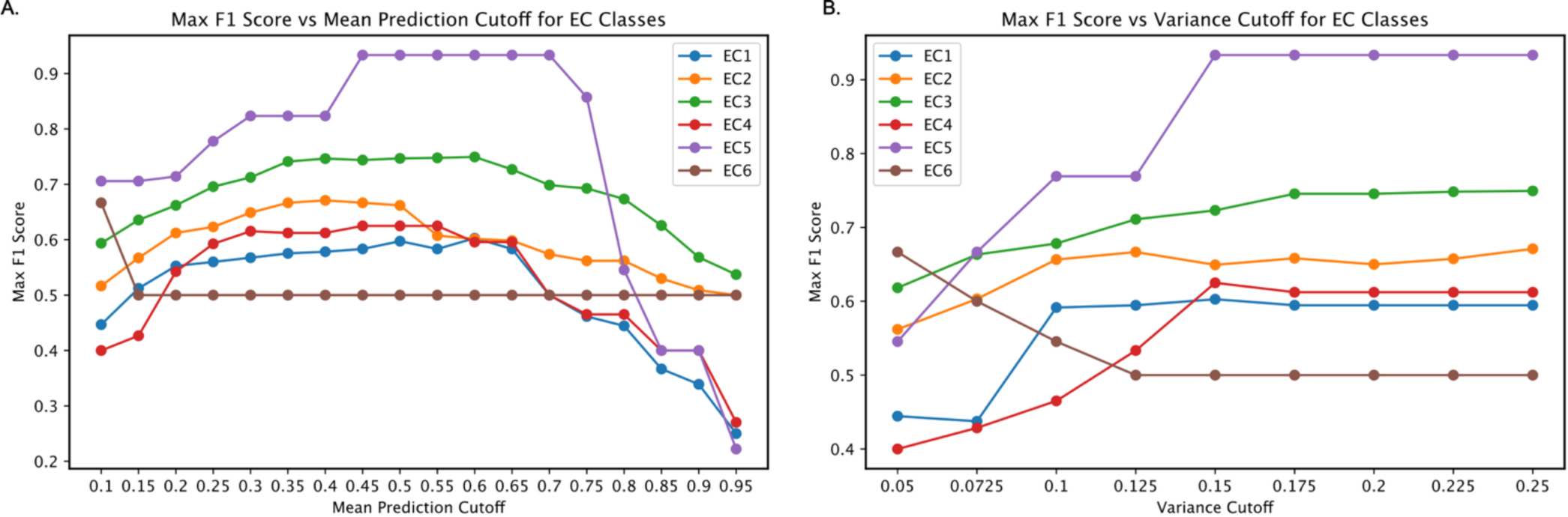
The maximum F1 score achievable with the Squidly ensemble model for each EC number in the CataloDB benchmark under varying mean prediction and variance thresholds. The default threshold of 0.6 and 0.225 determined using the uni3175 benchmark is relatively stable for each EC number. However, the limited data for EC 5 and 6 (N= 8, N=3, respectively) is such that users are recommended to further test the decision thresholds for their intended application.

**Figure S5:**
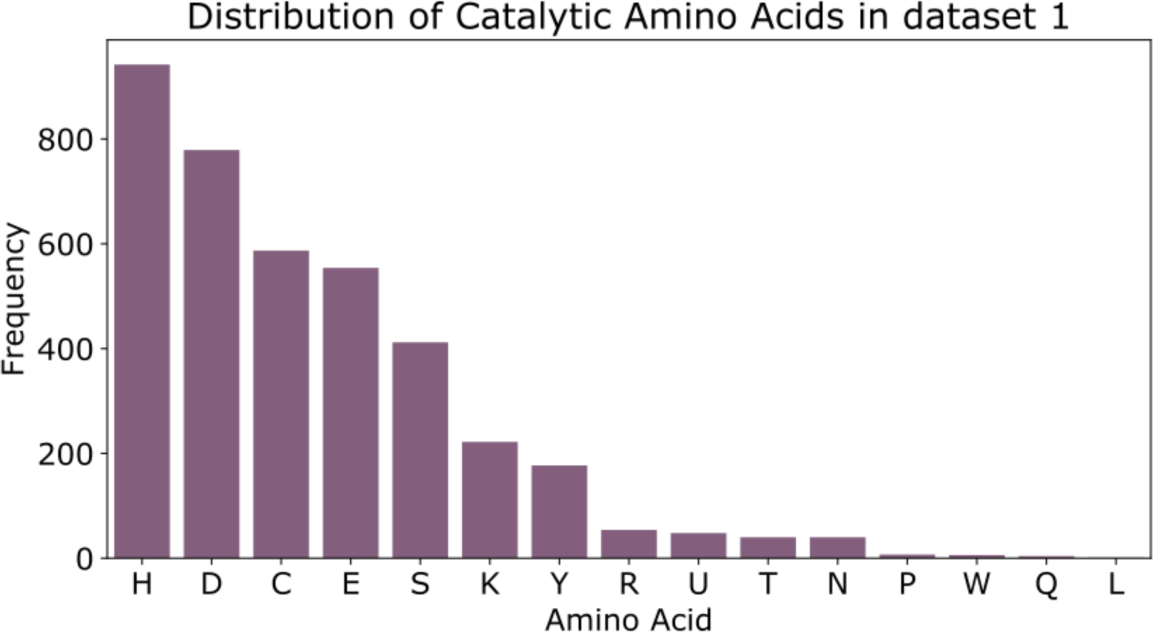
Amino acid distribution of Catalytic Residues in Dataset 1. Dataset 1 contains 3,873 catalytic residues, with 15 possible amino acids. The database is skewed towards certain amino acids which are commonly involved in catalysis.

**Figure S6:**
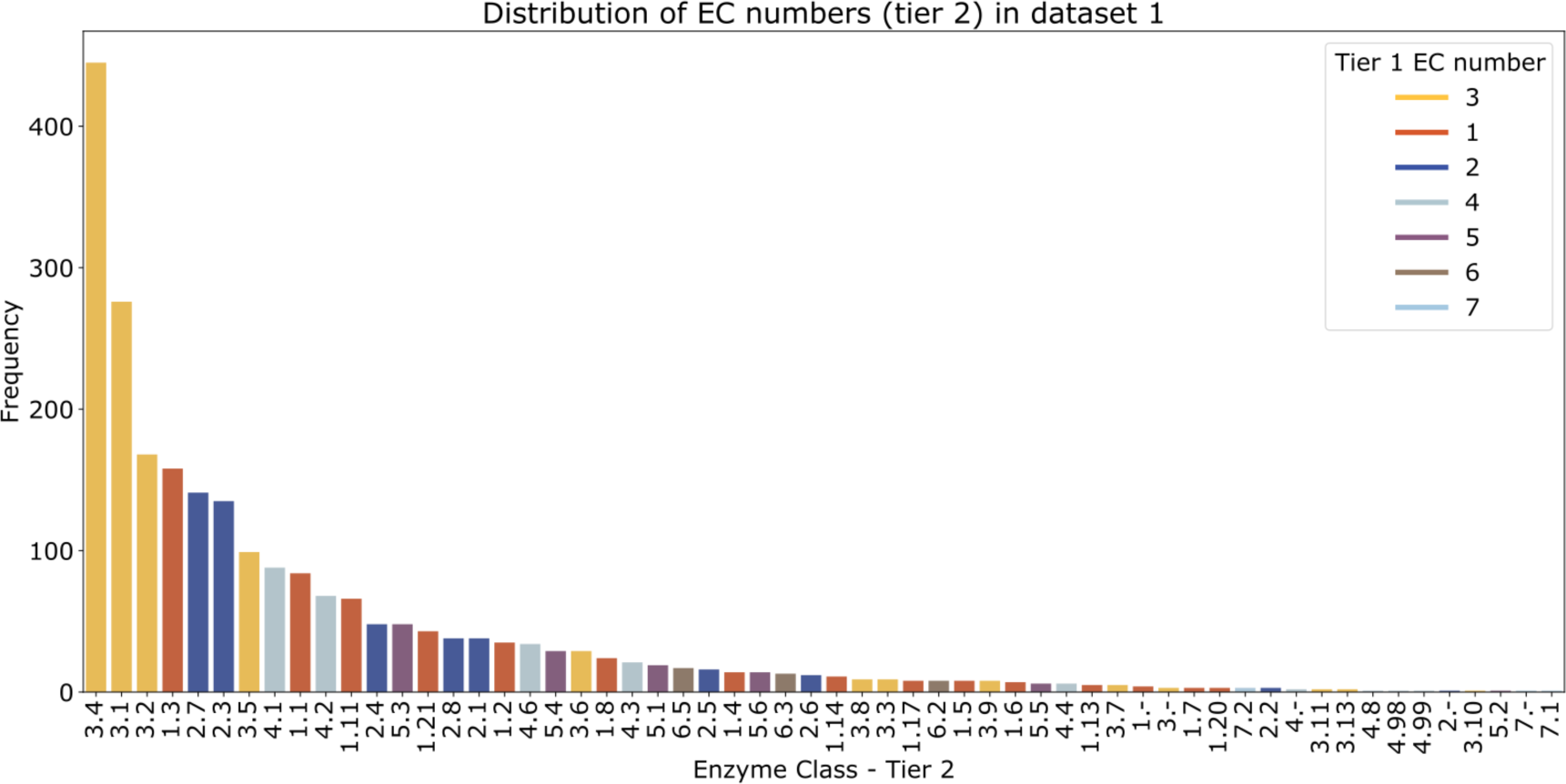
Dataset 1 EC Distribution. Dataset 1 is made up of 2,210 sequences from Swissprot. The sequences have experimental validation for the catalytic residue annotations. Validation sets are derived exclusively from this distribution. A clear bias exists for hydrolases (EC 3.X).

**Figure S7:**
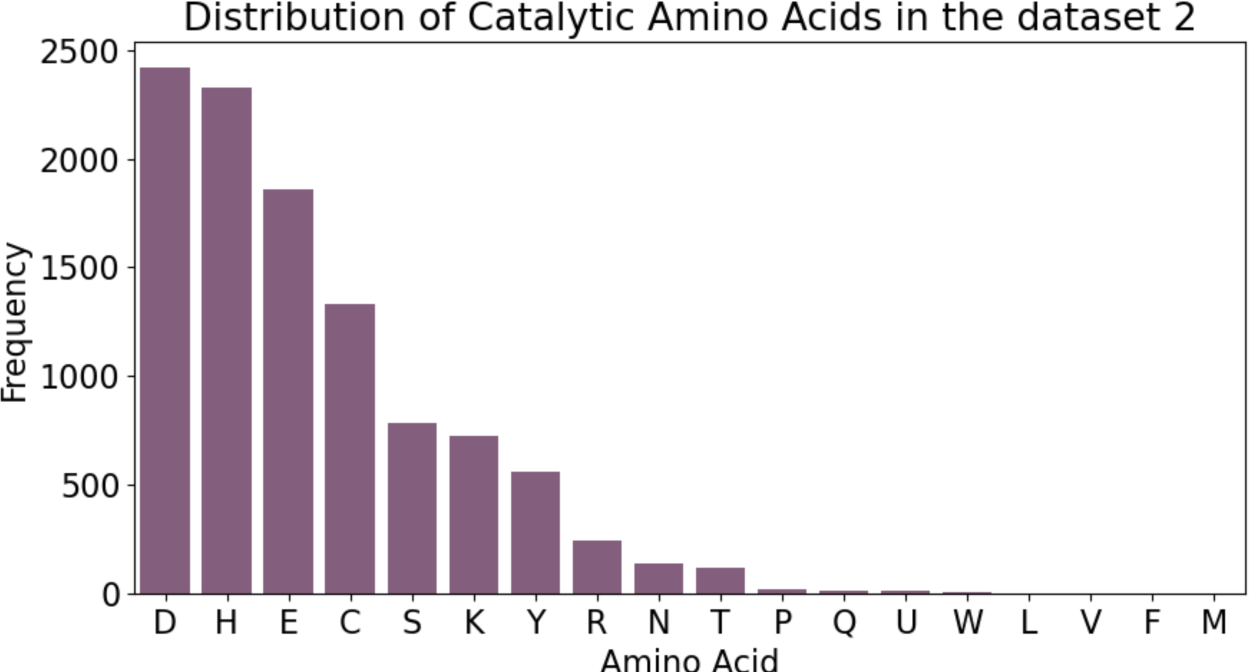
Amino acid distribution of Catalytic Residues in Dataset 2. Catalytic residue distribution. Dataset 2 contains 10,564 catalytic residues, with 18 possible amino acids. Amino acids M, V and F are additions not seen in dataset 1, although they exist in very few numbers.

**Figure S8:**
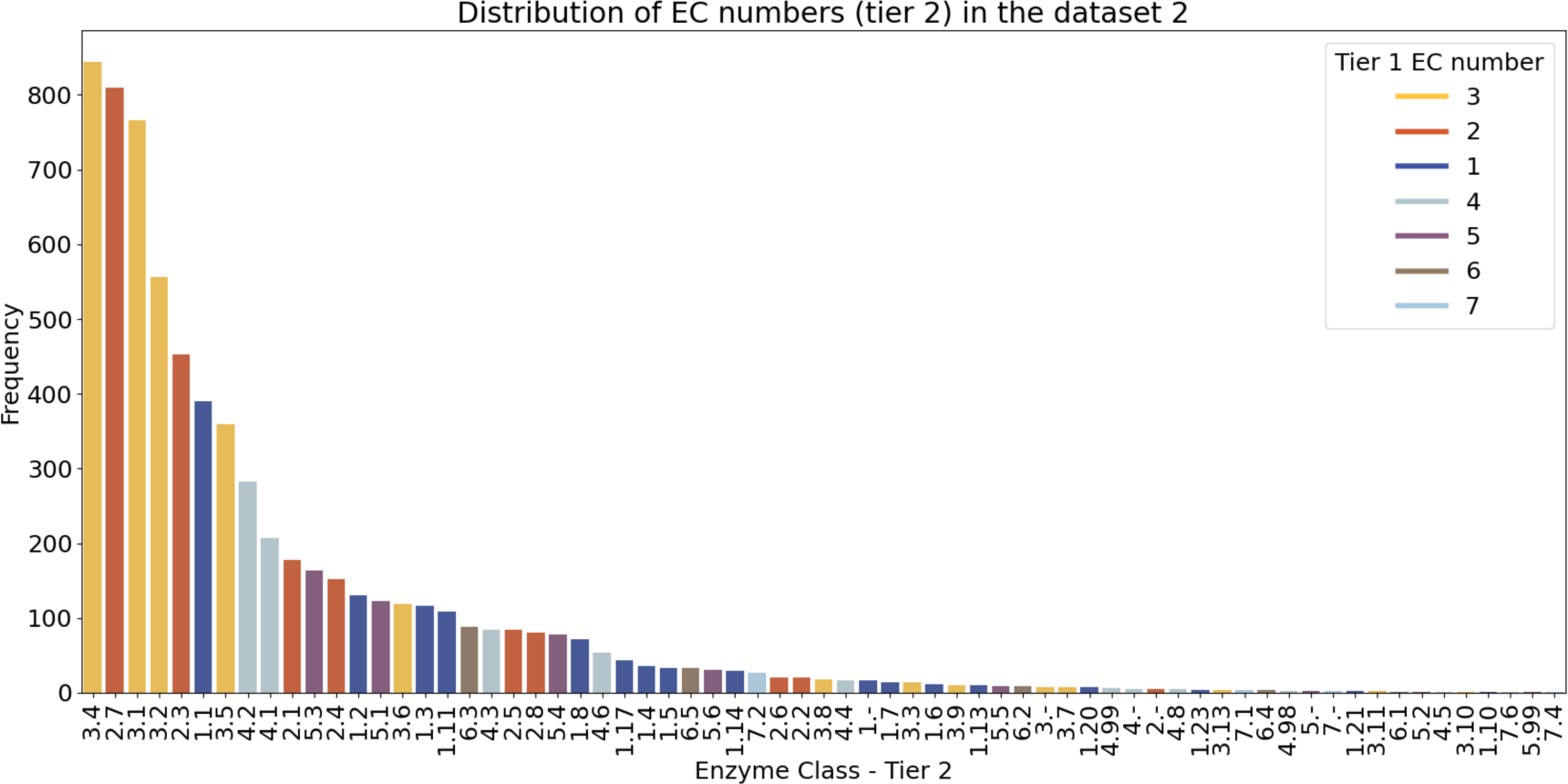
Dataset 3 EC Distribution. Dataset 3 is made up of 48,625 sequences. The sequences are taken from all the available proteins with catalytic site annotations in Swissprot. A very similar distribution is seen between datasets 3 and 2, with a notable increase in EC 2 again.

**Figure S9:**
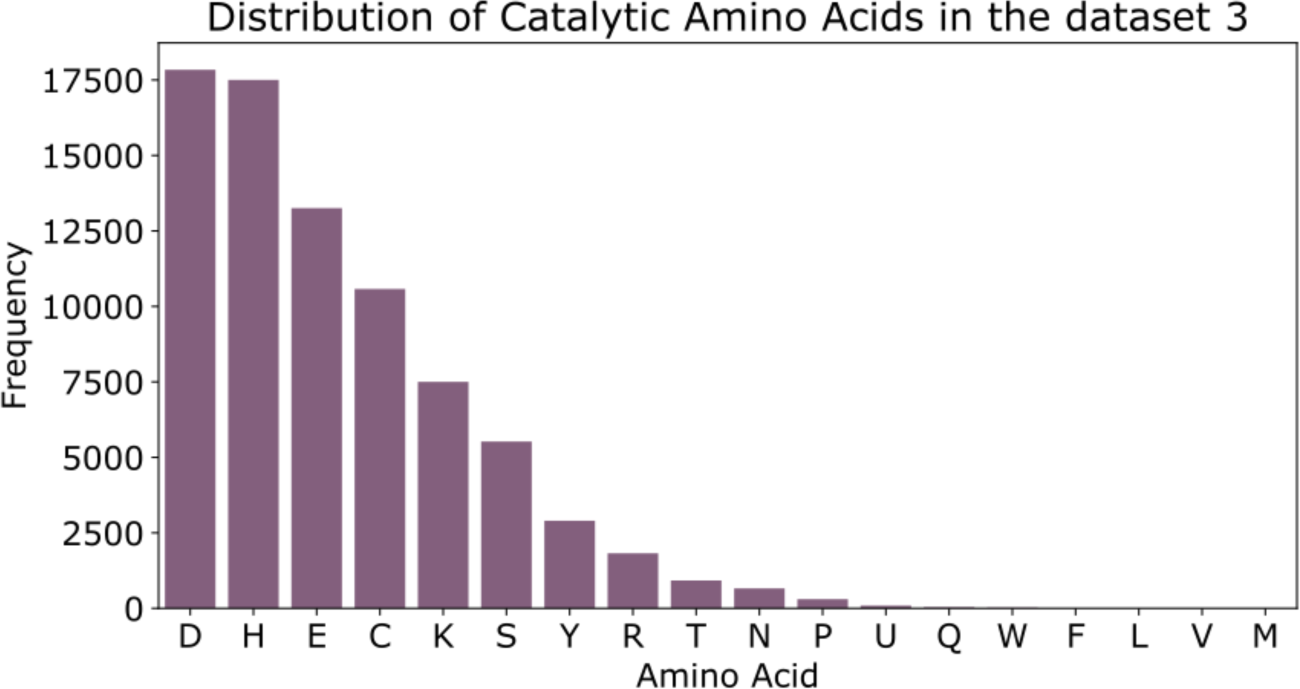
Amino acid distribution of Catalytic Residues in Dataset 3. Dataset 3 contains 78,966 catalytic residues, with 19 possible catalytic amino acids. Isoleucine is an additional amino acid not seen in dataset 1 or 2. The distribution seen here is very similar to that of dataset 1 and 2.

**Figure S10:**
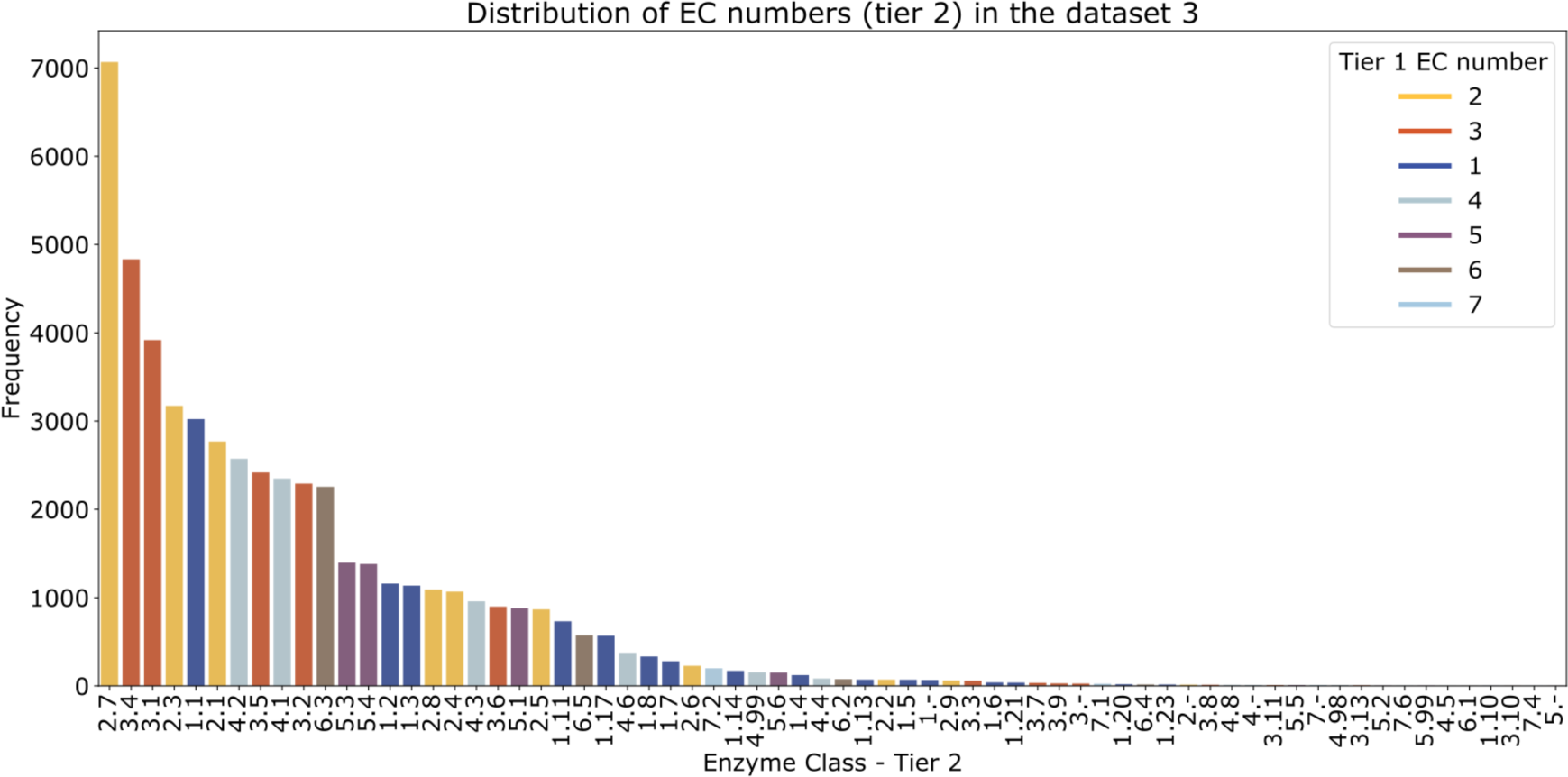
Dataset 3 EC Distribution. Dataset 3 is made up of 48,625 sequences. The sequences are taken from all the available proteins with catalytic site annotations in Swissprot. A very similar distribution is seen between datasets 3 and 2, with a notable increase in EC 2 again.

**Table S1.** Brief description of methods compared to Squidly in common benchmarks.

| **Study** | **Input** | **Method** |
| --- | --- | --- |
| **SCREEN**<sup>9</sup> | Structure, sequence embeddings, and evolutionary conservation. | Graph Neural Network which employs contrastive learning on EC numbers. |
| **AEGAN**<sup>10</sup> | Structure, Atchley Factors, PSSM conservation scores | Integrated graphical representations are generated using local protein networks and an amino acid spatial model with conservation scores and Atchley Factors included. A Graph Neural Network is trained to predict Catalytic residues. |
| **PREvail**<sup>17</sup> | Structure and amino acid sequence | Generates a range of sequence- and structure-based features representing conservation, structural descriptions, and network properties of the enzyme, and then performs feature selection and trains Random Forest models for prediction. |
| **CRpred**<sup>8</sup> | Sequence | Generates and predicts sequence-based features, including PSSM conservation scores and trains an SVM for prediction. |

**Table S2.** Benchmark datasets in prior work.

| **Dataset** | **Year of publication** | **Size (before sub-setting)** | **SCREEN Subset** | **SCREEN >90% identity** | **AEGAN Subset** | **AEGAN >90% identity** |
| --- | --- | --- | --- | --- | --- | --- |
| **EF family** <sup>35</sup> | 2007 | 293 | NA | NA | 196 | 3.6% |
| **EF superfamily** <sup>35</sup> | 2007 | 189 | 49 | 14.3% | 93 | 2.2% |
| **EF fold** <sup>35</sup> | 2007 | 236 | 60 | 10.0% | 123 | 3.3% |
| **HA superfamily** <sup>36</sup> | 2007 | 282 | 191 | 77.0% | 152 | 2.6% |
| **NN** <sup>37</sup> | 2002 | 163 | 95 | 83.1% | 108 | 2.7% |
| **PC** <sup>38</sup> | 2006 | 81 | 48 | 85.4% | 55 | 1.8% |
| **SCREEN training** <sup>9</sup> | 2024 | 1055 | 1055 | NA | NA | NA |
| **AEGAN training (Uni14230)** <sup>10</sup> | 2023 | 8435 | NA | NA | 8435 | NA |
| **AEGAN test (Uni3175)** | 2023 | 1955 | NA | NA | 1955 | 0.00% |

